# A CRISPR-based *Xenopus tropicalis* model for retroperitoneal liposarcoma with genetic control over the dedifferentiation process

**DOI:** 10.64898/2026.03.26.714450

**Authors:** Marthe Boelens, Dieter Tulkens, Arne Christiaens, Wout Houbart, Suzan Demuynck, David Creytens, Kris Vleminckx

**Author notes:** These authors contributed equally (Marthe Boelens, Dieter Tulkens).

## Abstract

Well- and dedifferentiated liposarcomas (WDLPS and DDLPS) are characterized by extensive copy-number alterations rather than recurrent gene-inactivating mutations, obscuring the molecular mechanisms that drive disease progression and, critically, the transition from well-differentiated to the more aggressive dedifferentiated tumor states. Despite marked differences in clinical behavior and prognosis, the regulatory events underlying adipocytic lineage destabilization in DDLPS remain poorly understood.

Here, we establish an *in vivo* model of retroperitoneal liposarcoma in *Xenopus tropicalis* through early embryonic mosaic perturbation of p53 and Rb pathway components. Combined disruption reproducibly induced retroperitoneal WDLPS development, demonstrating that pathway-level deregulation of the MDM2–p53 and CDK4–Rb axes is sufficient to initiate liposarcoma development *in vivo*.

Strikingly, additional perturbation of transcriptional co-activator *ep300* in this context resulted in increased tumor dedifferentiation, yielding lesions composed of spatially coexisting well- and dedifferentiated adipocytic states. In contrast, direct targeted disruption of downstream adipogenic regulators recurrently lost in human DDLPS, including *cebpa*, *g0s2*, and *dgat2*, failed to induce dedifferentiation in the same genetic context *in vivo*. These findings indicate that dedifferentiation cannot be explained by loss of downstream adipocytic effectors alone but instead reflects destabilization of higher-order regulatory programs governing adipocytic identity.

Together, these results establish an *in vivo* model that closely reflects the clinical situation on a pathway level and provides initial mechanistic insight into how adipocytic differentiation may become destabilized during disease progression. This framework offers a foundation for future studies leveraging higher-order and multi-omic approaches to dissect the molecular processes underlying the WDLPS-to-DDLPS transition.

## Introduction

Large-scale cancer genome sequencing studies have revealed that many malignancies are dominated by extensive copy number alterations (CNAs) and structural chromosomal rearrangements rather than recurrent gene-level mutations^1,2^. This is particularly evident in soft tissue sarcomas (STS), where complex-genome subtypes exhibit widespread chromosomal gains and losses affecting approximately 30–60% of the genome, often encompassing dozens of large-scale CNA events per tumor, and where overall CNA burden correlates with histological subtype and clinical outcome^3,4^. Although CNA landscapes have been extensively described and strongly associated with tumor behavior, they encompass large genomic regions containing dozens to hundreds of genes, making it difficult to pinpoint the specific genetic insults that drive tumor initiation (*i.e.* the inactivation of tumor suppressor genes or activation of oncogenes), progression, or dedifferentiation. As a result, despite strong genomic evidence implicating CNAs in sarcoma biology, the causal roles of individual genes within these altered regions remain poorly understood, creating a major bottleneck in translating CNA maps into actionable biological drivers.

Retroperitoneal liposarcoma (LPS), the most common soft tissue sarcoma in adults, represents a clear example of this limitation. Well-differentiated and dedifferentiated liposarcomas (WDLPS and DDLPS; which together represent 20% of all STS and approximately 60% of all LPS) are characterized by recurrent CNAs, most notably the defining amplification of the 12q13–15 region containing *MDM2* and *CDK4*^4^. This amplicon, typically located on supernumerary ring or giant marker chromosomes, spans a variable genomic segment containing numerous co-amplified genes within 12q13–15, including *HMGA2*, *TSPAN31*, *YEATS4*, *FRS2*, *GLI1* among other candidate oncogenes. In addition to amplification-driven oncogene activation, the 12q13–15 locus frequently undergoes chromosomal breakpoints that disrupt HMGA2, generating truncated transcripts or fusion genes that contribute to tumorigenesis^5^. Additional co-amplifications in distinct genomic regions, in particular the *JUN*-containing 1p32 locus, are frequently observed. Several of these genes have been implicated in tumor progression through incompletely understood mechanisms^6,7^.

While both WDLPS and DDLPS share an amplification-driven oncogenic backbone, progression to dedifferentiated disease is often accompanied by a substantial increase in genomic complexity. In addition to recurrent DDLPS-enriched gains on 5p and 14q, the dedifferentiated state can be associated with the acquisition of widespread additional copy-number losses^3,8–10^. Several additional loci, including 11q23–24, 19q13, 3q29, 9p22–24, and 17q21, have been linked to adverse clinical outcome. This increase in genomic complexity is possibly a consequence of destabilization of p53 protein by high levels of MDM2, which could promote large-scale genomic disruption events such as chromothripsis^9^, indicating that many of these CNAs may arise as byproducts of genomic instability rather than bona fide driver alterations. Given their presumed origin from a common mesenchymal progenitor^11^, progression toward dedifferentiation likely arises from secondary alterations emerging within a background of p53-associated genomic instability, with only a fraction contributing as true drivers of disease progression.

Because many of the genomic losses observed in DDLPS affect broad chromosomal regions, a central unresolved question is how these alterations translate into loss of adipocytic identity and emergence of dedifferentiated tumor states. Dedifferentiation is fundamentally a cell-state transition. Under physiological conditions, adipocytic lineage identity is maintained by a coordinated transcriptional program centered on regulators such as PPARG and CEBPA that enforce and stabilize terminal differentiation. In DDLPS, multiple components of this program are recurrently attenuated through genomic loss or epigenetic silencing, linking impaired adipocytic differentiation to aggressive disease behavior^12,13^.

To date, the biology of lineage destabilization is only partially reflected in current therapeutic strategies, which largely focus on shared amplification events rather than differentiation state control. Consequently, molecular treatment efforts in DDLPS have focused on targeting *MDM2* and *CDK4* amplification, with only limited clinical benefit^14^. This likely reflects the continued clinical management of well-differentiated and dedifferentiated liposarcoma as a single disease entity, resulting in suboptimal stratification of patients presenting with high-grade DDLPS and increased relapse rates. Distinct treatment approaches are therefore required, with a key focus on DDLPS disease management, as WDLPS patients have a substantially more favorable prognosis, with reported 5-year survival rates of 82% versus 48% and a median survival time of 16.2 years versus 4.5 years, respectively^15^.

Identification and *in vivo* functional validation of novel targetable dependency genes is thus needed to guide and accelerate molecular treatment strategies for the more aggressive and therapy-resistant dedifferentiated state. Achieving a comprehensive understanding of DDLPS biology requires omics-based dissection of tumors in their native context. Although analyses of surgical specimens have yielded important insights^10,16,17^, progress is constrained by limited, patient-specific material typically obtained from partial resections or biopsies that incompletely capture tumor heterogeneity. This underscores the need for robust *in vivo* model systems to systematically resolve candidate driver and dependency genes beyond what can be achieved with available cell line and PDX platforms. While genetically engineered mouse and zebrafish models have yielded valuable biological insights, they are typically constrained in scalability for systematic driver and dependency gene identification and often rely on lineage-restricted targeting strategies, which can limit their ability to model the cellular plasticity and molecular heterogeneity of WD/DDLPS^18,19^.

Here, we employ *Xenopus tropicalis* as a rapid and scalable *in vivo* system to functionally assess candidate drivers and dependencies of retroperitoneal liposarcoma. Using multiplex CRISPR/Cas9- mediated genome editing, we establish a liposarcoma model that recapitulates key molecular, histopathological, and pathway-level features of the human disease, including WDLPS to DDLPS transition. Importantly, targeted genome editing in specific blastomeres at early embryonic stages, enabling lineage enrichment rather than strict lineage restriction, increasing the likelihood that oncogenic perturbations arise within the appropriate cell-of-origin^20^. By enabling direct evaluation of (epi)-genetic events driving liposarcoma progression within a diploid vertebrate organism, this work addresses a key gap between molecular research and clinically relevant tumor biology.

## Results

### Multiplex disruption of p53 and Rb pathways generates high-penetrance retroperitoneal liposarcoma in *Xenopus tropicalis*

Building on our longstanding interest in Rb pathway-driven malignancies (including retinoblastoma, pancreatic neuroendocrine carcinoma, choroid plexus carcinoma, and high-grade glioblastoma^21,22^), we employed multiplex CRISPR/Cas9-mediated genome editing in *Xenopus tropicalis* to achieve combined inactivation of the p53 and Rb tumor suppressor pathways^23,24^. Targeting *tp53* together with all three Rb family members (*rb1, rbl1,* and *rbl2*) enforces complete pathway-level disruption. This functionally recapitulates the core oncogenic mechanism underlying human well-differentiated and dedifferentiated liposarcoma, where amplification of the 12q13–15 locus, encompassing *MDM2* and *CDK4*, leads to suppression of p53 activity and inactivation of Rb proteins, resulting in uncontrolled cell proliferation. Although mediated by distinct genomic alterations, both contexts converge on effective loss of p53- and Rb-dependent tumor suppressive control.

To additionally model the differentiation plasticity characteristic of dedifferentiated disease, we incorporated targeting of *ep300*, a transcriptional co-activator and chromatin regulator essential for maintaining lineage-specific gene expression programs^25^. In human liposarcoma, progression from well-differentiated to dedifferentiated tumors is found to be frequently associated with accumulation of additional copy-number losses that preferentially affect differentiation-stabilizing and chromatin-regulatory loci^8^. Functional ablation of *ep300* was therefore designed to approximate the consequences of these dedifferentiation-associated losses at the pathway level.

Strikingly, combined CRISPR-mediated targeting of *tp53*, all *rb* gene family members, and *ep300* in early *Xenopus* embryos, resulted in mosaic mutant animals (hereafter referred to as the **Δp53/Rb/p300** cohort) that showed rapid and highly penetrant formation of retroperitoneal liposarcoma (from 70 days post fertilization (dpf) onwards with 41.7% incidence, n=24), amongst well-established other tumor types such as retinoblastoma and scPaNEC^21,22^ (Fig. 1A-C, Supplementary Table 1A). Macroscopic examination revealed opaque adipose strands with irregular, finger-like extensions and increased vascularization, in contrast to intra-animal control fat, which appeared yellow and glossy, reflecting abundant lipid content (Fig. 1D). Histological analysis based on hematoxylin and eosin (H&E) stained tissue samples revealed that individual tumors frequently (*i.e.* 70%, n=10) contained both well-differentiated and dedifferentiated components, with regions displaying recognizable adipocytic morphology characteristic of well-differentiated liposarcoma interspersed with areas of increased cellularity, nuclear atypia, and loss of adipocytic features consistent with dedifferentiated disease (Fig. 1E). These findings establish a genetically defined *Xenopus tropicalis* model that captures both the core oncogenic drivers and the differentiation plasticity characteristic of this disease. This co-existence of distinct differentiation states that typify clinical WD/DDLPS, provides a unique *in vivo* framework to directly compare molecular features underlying liposarcoma progression and dedifferentiation.

**Figure 1.**
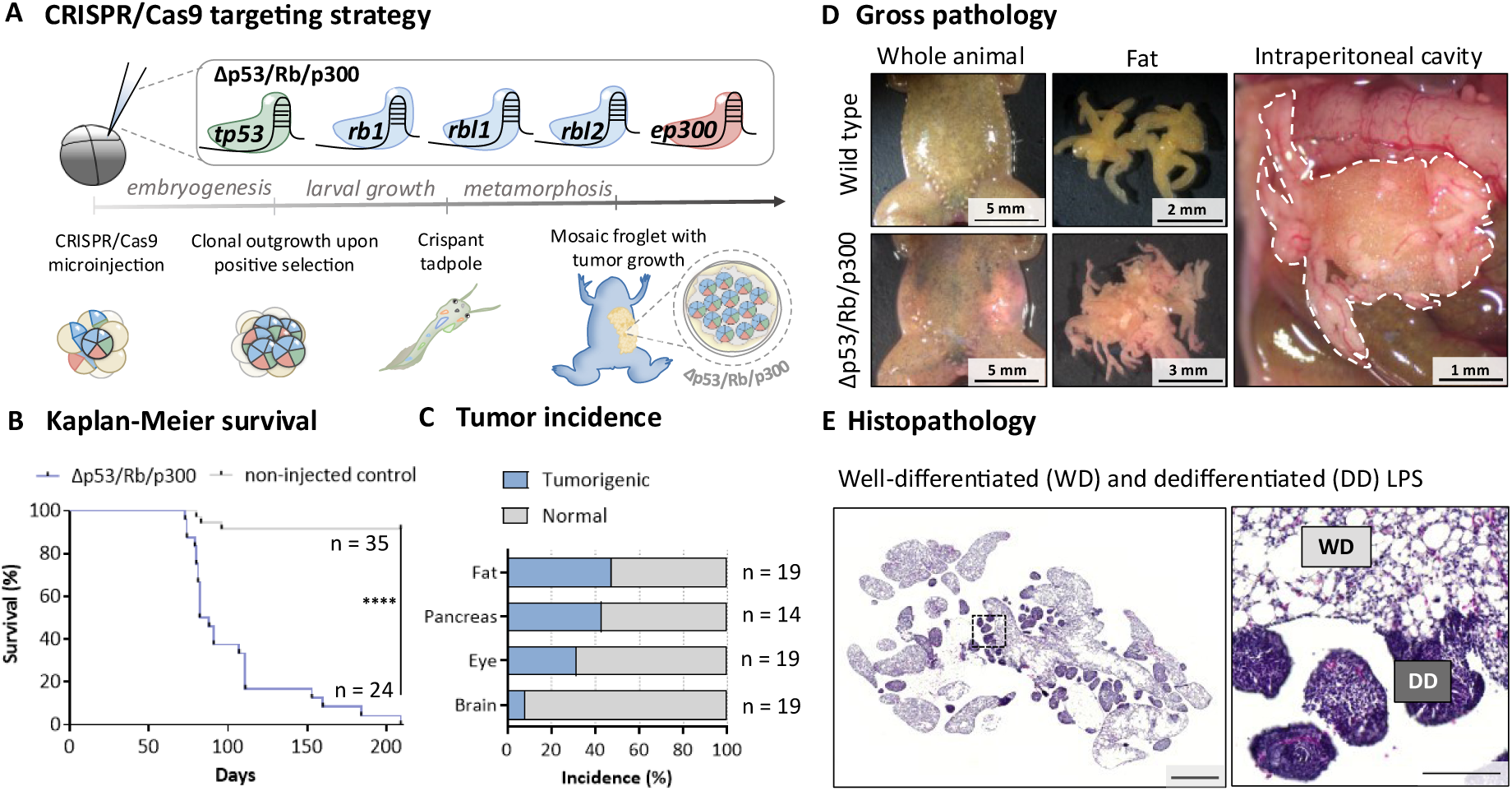
CRISPR/Cas9-mediated multiplexed knockout of tp53, all rb family members and ep300 induces liposarcomas in Xenopus tropicalis. **(A)** CRISPR/Cas9 targeting strategy for tp53, rb1, rbl1, rbl2 and ep300. Microinjection of precomplexed Cas9 and guide RNAs simultaneously targeting all five genes generates somatic mosaicism. Positive selection during development results in mosaic mutant froglets that develop tumors within 100 days. **(B)** Kaplan–Meier survival curves show a significant decrease in survival in injected versus non-injected cohorts. Δp53/Rb/p300 animals (n = 24) versus controls (n = 35). ****P < 0.0001. **(C)** Tumor incidence by anatomical site with fat tumors being highly penetrant. n indicates animals examined. **(D)** Gross pathology of representative normal fat from wild-type animals (top) and liposarcoma tissue from Δp53/Rb/p300 animals (bottom). White dashed outline indicates the retroperitoneal liposarcoma in the peritoneal cavity. **(E)** H&E histology showing well-differentiated (WD) and dedifferentiated (DD) liposarcoma regions. Scale bars: 500 μm and 100 μm (inset).

### *Xenopus tropicalis* liposarcoma cells show engraftment potential upon transplantation

To assess the malignant potential and proliferative capacity of Δp53/Rb/p300-derived liposarcomas, we performed transplantation experiments using a recently established *Xenopus tropicalis rag2*^−/−^ line, which lacks mature T and B lymphocytes and permits stable engraftment of transplanted tumor cells^26^.

Purified single-cell suspensions derived from primary liposarcoma donor tissue were injected intraperitoneally (IP) and subcutaneously (SC, dorsal region) into an adult *rag2*^−/−^ recipient animal (Fig. 2A). The host developed overt signs of disease 51 days following transplantation and was sacrificed at this time point. Post-mortem analysis revealed robust tumor engraftment within the intraperitoneal cavity and lungs, with histological features closely resembling those of the original donor liposarcoma, and positive staining for the proliferation markers PHH3 and PCNA, confirming the proliferative nature of the tumor grafts (Fig. 2B and Supplementary Fig. 1). In addition, clear subcutaneous tumor growth was observed at the dorsal injection site, confirming efficient engraftment across multiple anatomical locations. We independently confirmed engraftment of liposarcoma cells from the same Δp53/Rb/p300 donor in a parallel transplantation experiment using a second rag2^−/−^ recipient. A prominent liposarcoma graft formed at the injection site, with additional smaller lesions detected in the lungs, further indicating robust engraftment (Supplementary Fig. 2).

**Figure 2.**
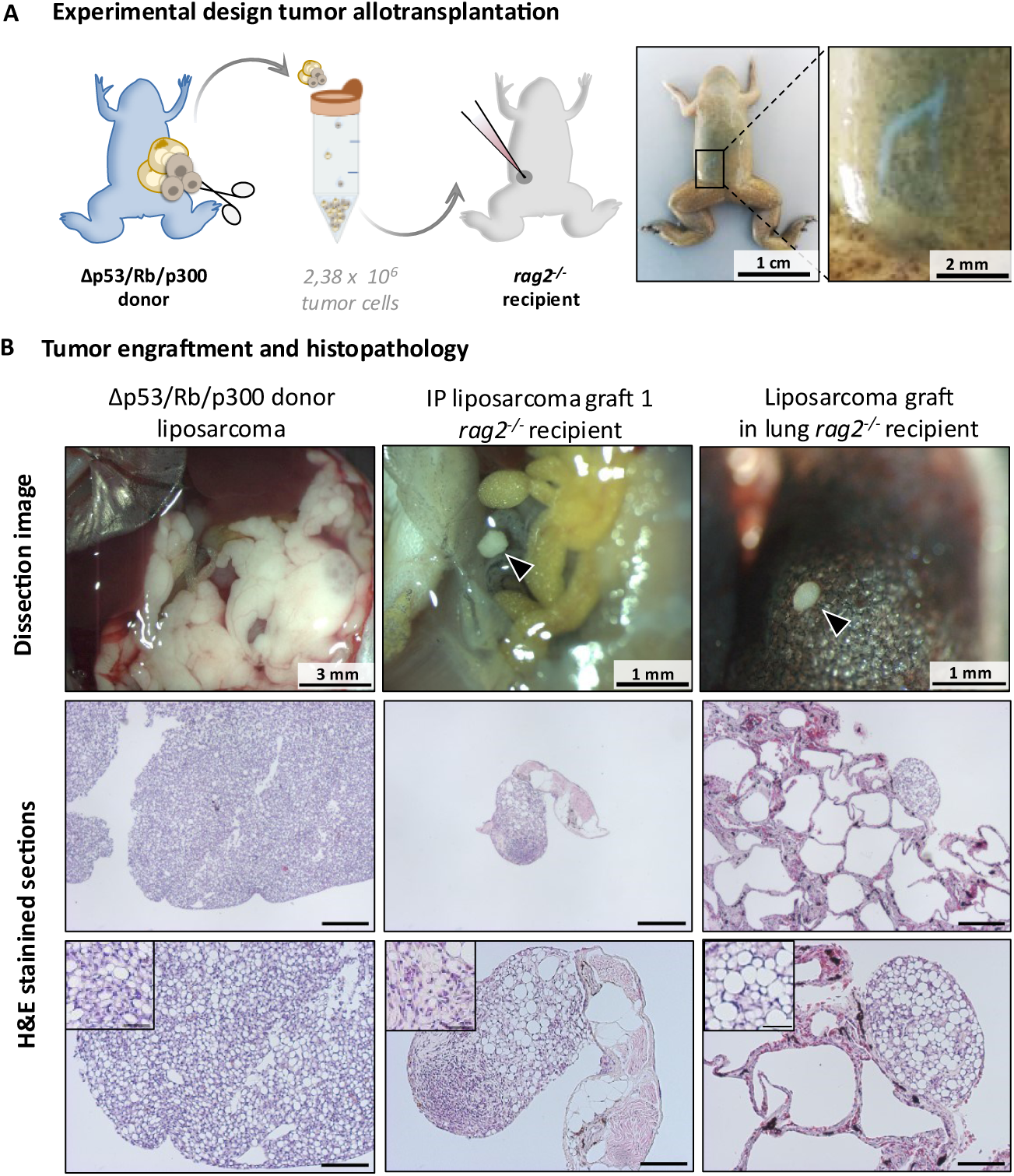
Allotransplantation of Δp53/Rb/p300 liposarcomas in immunodeficient rag2^−/−^ Xenopus tropicalis. **(A)** Experimental workflow. Liposarcomas harboring Δp53/Rb/p300 mutations were dissected from donor frogs, dissociated, and 2.38 × 10⁶ cells transplanted intraperitoneally into rag2⁻/⁻ immunodeficient recipients. Representative images show a recipient frog at 4 weeks post-transplantation with a visible scar at the injection site (inset shows magnified view of the scar). **(B)** Pathological characterization of tumor engraftment. Top row: gross pathology of primary donor tumor (left), intraperitoneal graft in recipient (middle, black arrow head), and pulmonary graft (right, black arrow head). Middle and bottom rows: H&E-stained sections at low and high magnification showing preserved liposarcoma histology with lipoblasts (cells with cytoplasmic lipid vacuoles and peripheral nuclei, see insets). Note dense cellular regions intermixed with adipocyte-like areas characteristic of well-differentiated liposarcoma. Scale bars: 250 μm (middle row), 100 μm (bottom row), 25 μm (inset).

Collectively, these transplantation experiments demonstrate that Δp53/Rb/p300-derived liposarcomas are highly tumorigenic, capable of engrafting at multiple anatomical sites, and retain histological identity following primary transplantation.

### Genetic profiling reveals *tp53*, *rb1* and *rbl1* as core driver alterations underlying liposarcoma development

To characterize the genetic alterations underlying tumor formation, we performed deep amplicon sequencing of all five targeted loci in eight independent Δp53/Rb/p300-derived liposarcoma samples as well as in tumors derived from subsequent injection cohorts with comparable targeting of the intended loci, and analyzed the genome editing events using the BATCH-GE analysis tool (Fig. 3A-B and Supplementary Fig. 3 and 4; Supplementary Table 1B)^27^. Deep sequencing of CRISPR target sites in embryo pools at 3 days post injection (dpi) revealed a heterogeneous spectrum of low-frequency indels characteristic of mosaic editing. In contrast, independent liposarcomas showed strong enrichment of dominant alleles at each targeted locus. For example, the most abundant *tp53* variant increased from 18.6% in embryo pools to 40–89% across tumors. Similar enrichment of dominant variants was observed for *rb1, rbl1, rbl2,* and *ep300*, indicating clonal expansion of cells carrying selected CRISPR-induced mutations during tumor development (Supplementary Fig. 3; Supplementary Table 1B).

**Figure 3.**
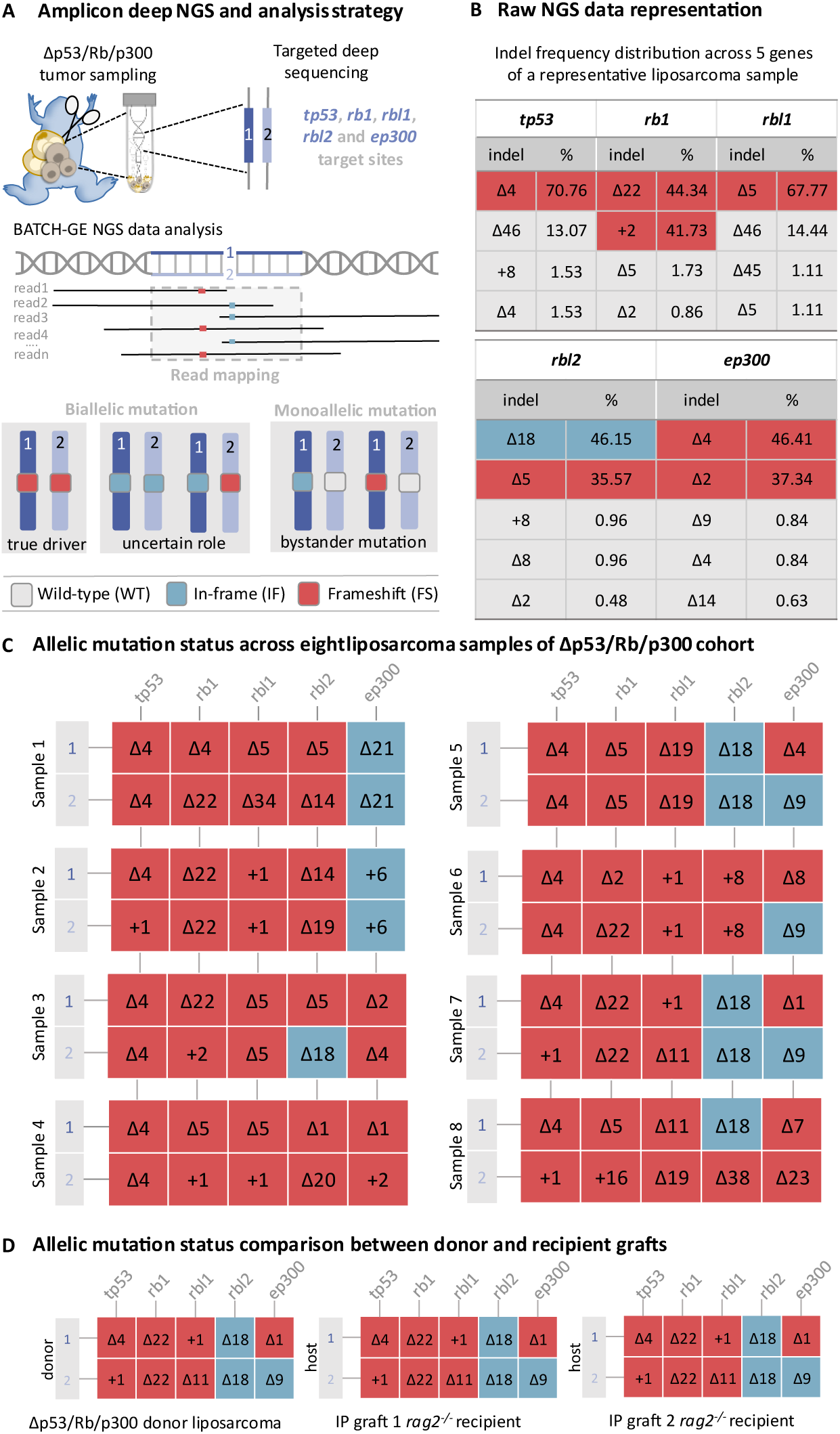
Deep amplicon sequencing reveals positive selection on biallelic mutated gene status in Δp53/Rb/p300 liposarcomas. **(A)** Targeted amplicon sequencing workflow. Xenopus tropicalis liposarcoma tumor DNA was PCR-amplified around the cut site of CRISPR-edited loci (tp53, rb1, rbl1, rbl2 and ep300), sequenced at high depth and analyzed using BATCH-GE to quantify indel frequencies and infer allelic mutation status. **(B)** Representative indel frequency distribution for a single tumor sample across five target genes. Indels are shown as percentage of total reads per locus. Dominant indel reads (>35%) are highlighted in red (frameshift) or in blue (in-frame). Indel reads with a value over 65% are interpreted as coming from the two alleles. **(C)** Allele-specific mutation status across eight tumors. Two rows per sample represent inferred alleles. **(D)** Comparison of allelic mutation status between a donor liposarcoma (i.e. Sample 7) and matched recipient grafts, demonstrating preservation of CRISPR-induced mutations and genetic clonal stability upon transplantation.

**Figure 4.**
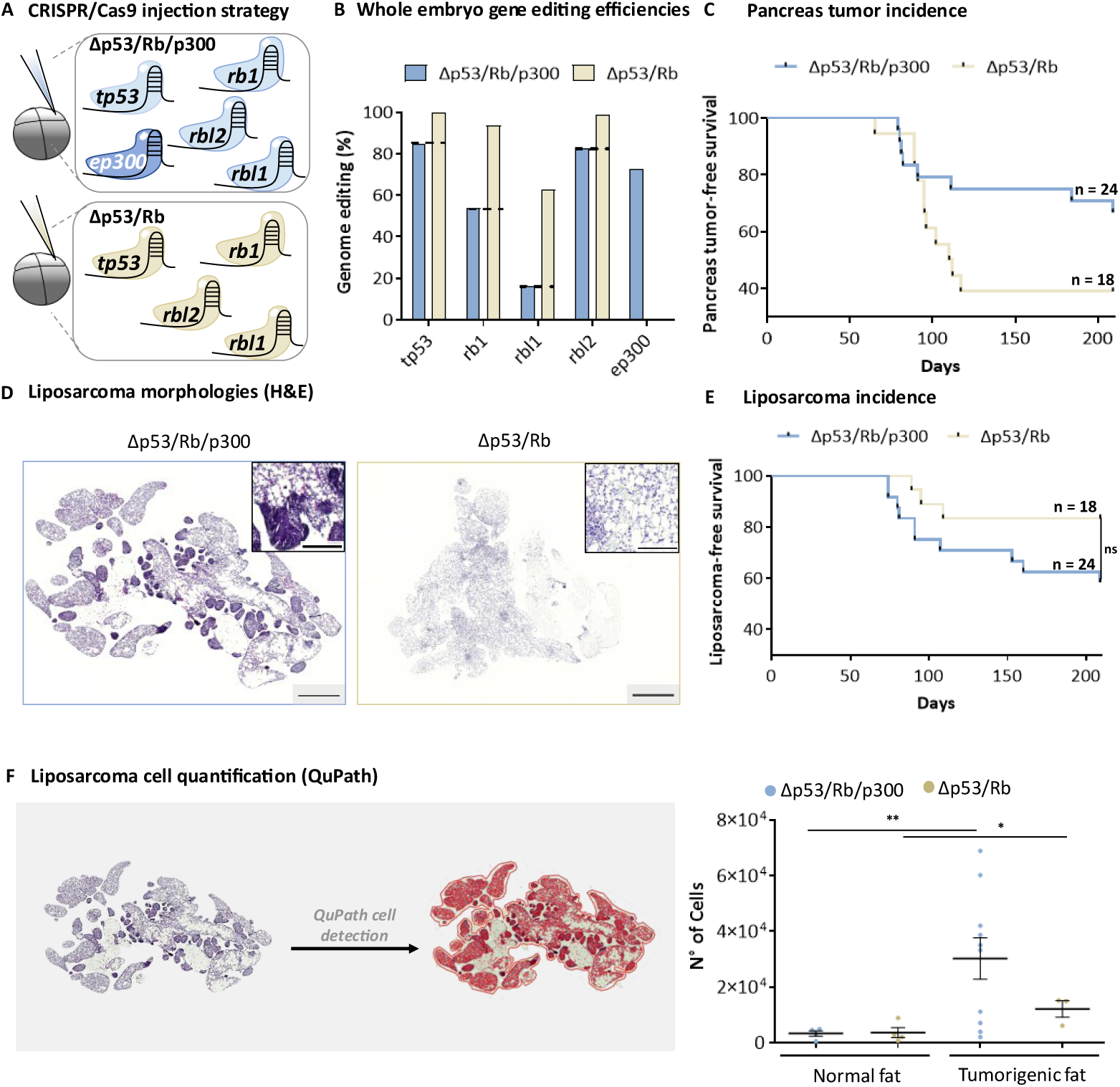
Loss of ep300 drives liposarcoma dedifferentiation. **(A)** Experimental design comparing multiplex CRISPR targeting of tp53 and rb family genes with or without ep300 disruption. **(B)** Calculated whole-embryo genome editing efficiencies for each targeted gene across injection cohorts. Total efficiencies were extrapolated by multiplying measured efficiencies by four, as embryos were injected in one of four blastomeres. Values exceeding 100%, likely due to incomplete cytokinesis at the time of injection allowing reagent diffusion to the sister blastomere, were normalized such that the most efficient sgRNA was set to 100%. **(C)** Pancreas tumor-free survival compared between Δp53/Rb/p300 and Δp53/Rb cohorts. **(D)** Representative H&E-stained sections illustrating liposarcoma morphology across Δp53/Rb/p300 and Δp53/Rb conditions, with mixed WD/DD regions and complete WD tumor tissue, respectively. **(F)** Quantification of liposarcoma cell numbers using QuPath. For each animal, the largest tumor section was analyzed (representative image shown on the left, all are listed in Supplementary Fig. 6). Four representative healthy littermate adipose tissue samples were included as controls to establish baseline adipocyte cellularity.

All eight liposarcoma samples exhibited extensive biallelic disruption across the five targeted genes, *tp53*, *rb1*, *rbl1*, *rbl2*, and *ep300*; with remarkable consistency in their mutational profiles (Fig. 3C and Supplementary Fig. 4; Supplementary Table 1B). Notably, *tp53*, *rb1*, and *rbl1* exclusively harbored biallelic frameshift mutations inducing indels, potentially generating premature stop-codons upstream of the final exon–exon junction, thereby likely targeting the mutant transcripts for nonsense-mediated mRNA decay (NMD)^28^. The uniform selection of such truncating alleles at these loci strongly suggests that functional inactivation of *tp53*, *rb1*, and *rbl1* confers a selective advantage during liposarcoma development, supporting their role as core tumor suppressors in this context.

By contrast, the mutational landscapes of *rbl2* and *ep300* were more heterogeneous. Although both genes were consistently disrupted on both alleles with clonal enrichment, multiple tumors retained in-frame indels, likely resulting in the loss or gain of amino acids (Fig.3C and Supplementary Fig. 4; Supplementary Table 1B highlighted)^29^. This pattern raises the question if, unlike *tp53*, *rb1*, and *rbl1*, functional variants of the Rbl2 and the p300 proteins are generated, and whether partial loss of function or hypomorphic alleles at these loci may be sufficient, or potentially advantageous, during liposarcoma evolution.

The same trend was observed after propagation of the tumor. To evaluate genetic stability following transplantation, two independent liposarcoma grafts floating freely within the intraperitoneal cavity of *rag2⁻/⁻* host were isolated and analyzed in parallel with a fragment of the original donor tumor by targeted amplicon deep sequencing. This analysis revealed complete concordance between donor and graft samples, with identical mutational variants detected across all targeted loci, demonstrating preservation of the monoclonal genetic architecture following transplantation (Fig. 3D).

### Loss of p300 in the context of disrupted p53 and Rb pathways contributes to liposarcoma dedifferentiation

Based on the mutational profiles observed in Δp53/Rb/p300 liposarcomas, we next sought to functionally assess the contribution of p300 and Rbl2 to liposarcoma development and differentiation state. To this end, we generated an additional multiplex CRISPR/Cas9 cohorts in which either the *ep300* of *rbl2* sgRNA was omitted (hereafter referred to as **Δp53/Rb** and the **Δp53/Rb^−Rbl2^ cohort**), while maintaining complete p53 and Rb pathway disruption (Fig. 4A and Supplementary Fig. 5A). To exclude confounding effects due to differences in genome editing efficiency, sgRNA concentrations were modestly increased in these cohorts. Whole-embryo amplicon sequencing confirmed that editing efficiencies at the remaining target loci exceeded those achieved in the Δp53/Rb/p300 condition (Fig. 4B and Supplementary Fig. 5B; Supplementary Table 1, Sheet A).

As a control readout, we first assessed pancreatic tumor incidence, which in this model depends on concurrent disruption of *rb1* and *rbl1*. Pancreatic tumors were most frequent in the Δp53/Rb cohort (61.1%, n = 18), followed by the Δp53/Rb^−Rbl2^ cohort (50.0%, n = 14), and least frequent in the Δp53/Rb/p300 cohort (33.3%, n = 24) (Fig. 4C and Supplementary Fig.5C, Supplementary Table 2, Sheet B and E). These frequencies closely matched the estimated probability of simultaneous *rb1/rbl1* editing inferred from whole-embryo sequencing (58.4%, 27.5%, and 8.8% for Δp53/Rb, Δp53/Rb^−Rbl2^, and Δp53/Rb/p300, respectively), supporting comparable genome editing performance across conditions.

**Figure 5.**
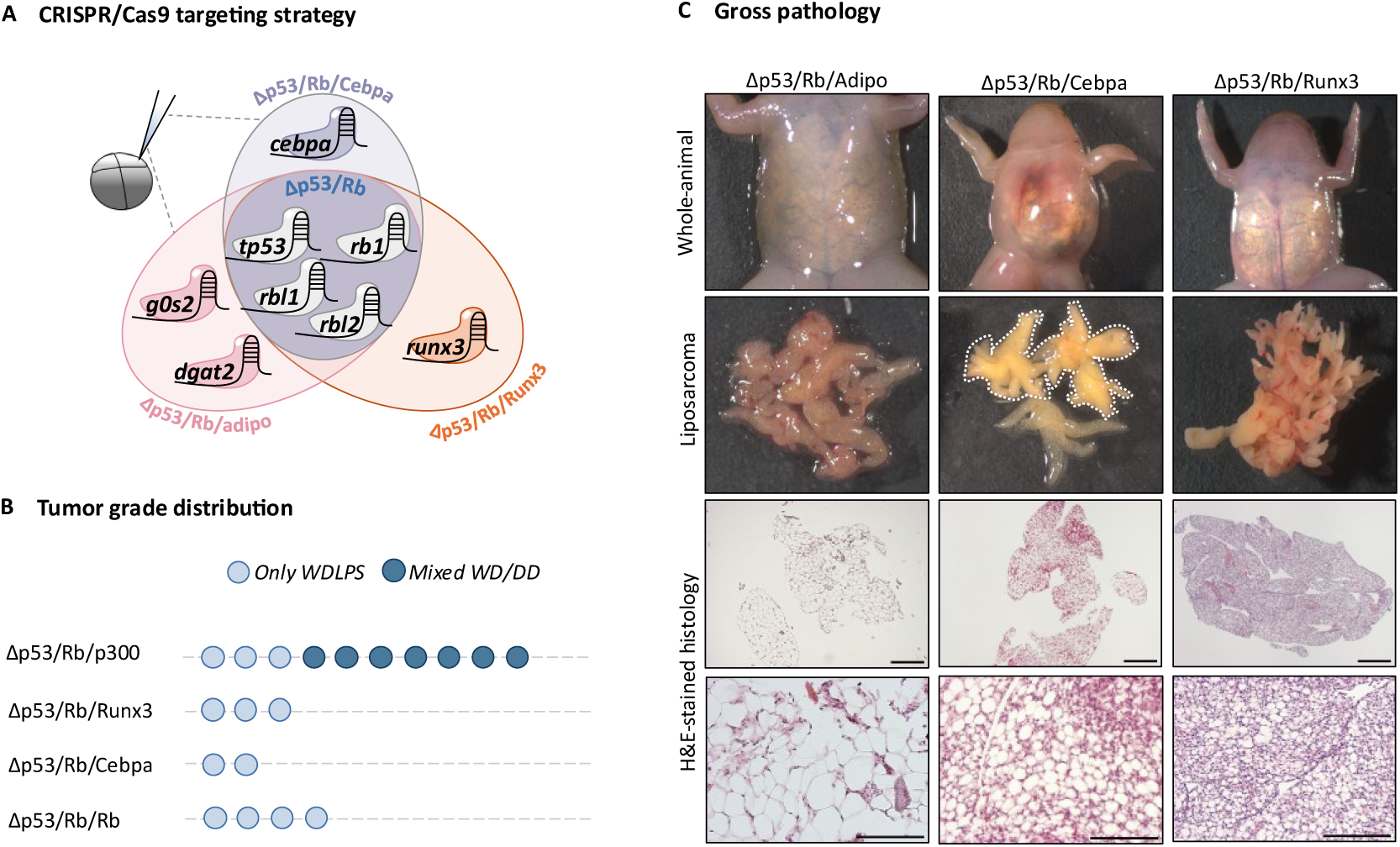
Direct disruption of adipogenesis effector genes is insufficient to drive dedifferentiation. **(A)** Schematic overview of the CRISPR/Cas9 multiplex targeting strategy used for identification of DDLPS candidate co-driver genes within a constant Δp53/Rb background. **(B)** Tumor grading analysis demonstrating absence of dedifferentiated liposarcoma in additional injection cohorts lacking ep300 targeting. Each circle represents one tumor sample; light blue indicates a well-differentiated liposarcoma (WDLPS) phenotype and dark blue a mixed well-differentiated/dedifferentiated (WD-DD) phenotype. **(C)** Gross pathological comparison of the three additional multiplex cohorts, with the highest-grade tumor per cohort shown as representative example.

Strikingly, retention of wild type p300 and Rbl2 was associated with a marked reduction in liposarcoma incidence. Whereas liposarcomas developed in 41.7% of Δp53/Rb/p300 animals (n = 24), incidence decreased to 16.7% in the Δp53/Rb cohort (n = 18, Mantel–Cox test, p = 0.0878; Fig. 4E; Supplementary Table 2, Sheet C), and even to 7.1% in the Δp53/Rb^−Rbl2^ cohort (n = 14, Mantel–Cox test, p < 0.05; Supplementary Fig. 5E; Supplementary Table 2, Sheet F). Thus, loss of *ep300* or *rbl2* substantially increases LPS tumor penetrance in the setting of p53 and Rb pathway disruption.

Histopathological evaluation further demonstrated that p300 loss profoundly influenced tumor differentiation state. Δp53/Rb/p300 liposarcomas frequently contained co-existing well-differentiated and dedifferentiated regions, whereas tumors arising in the Δp53/Rb cohort were uniformly well-differentiated and lacked overt dedifferentiated areas (Fig. 4D). Consistently, QuPath-based quantitative analysis demonstrated a substantially higher increase in cellularity in Δp53/Rb/p300 liposarcomas relative to normal adipose tissue (9.3-fold, p < 0.01), compared to Δp53/Rb tumors (3.3-fold, p < 0.05) (Fig. 4F and Supplementary Fig. 6; and Supplementary Table 2, Sheet D). Together, these data indicate that while combined p53 and Rb pathway disruption is sufficient to initiate well-differentiated liposarcoma, additional loss of p300 enhances tumor cellularity and promotes progression toward a less differentiated state.

The Δp53/Rb^−Rbl2^ cohort also showed a reduction in overall tumor cellularity relative to the Δp53/Rb/p300 cohort (Supplementary Fig. 5F and 6; Supplementary Table 2, Sheet G), however, the tumor histology remained unchanged (Supplementary Fig. 5D). These findings indicate that *rbl2* disruption is not strictly required as a co-driver of tumor initiation or morphological progression along the WDLPS–DDLPS spectrum. Instead, the phenotypic differences likely reflect context-dependent functional redundancy within the Rb family during early embryonic multiplex targeting^30–32^, whereby simultaneous perturbation of multiple pocket proteins, even in the presence of in-frame alleles, may suffice to compromise compensatory cell cycle control.

Together, these data identify p300 as a key modulator of liposarcoma penetrance and malignancy, reflected by increased tumor cellularity and progression toward a less differentiated state in the Δp53/Rb/p300 model. While combined disruption of the p53 and Rb pathways is sufficient to initiate well-differentiated liposarcoma, the context-dependent functional redundancy within the Rb family likely permits tumor initiation following multiplex targeting in early embryonic progenitors, even in the presence of partially functional in-frame alleles. In contrast, additional loss of p300 promotes increased tumor cellularity and the emergence of dedifferentiated regions, consistent with a role for transcriptional co-activation and chromatin regulation in maintaining adipocytic tumor identity.

### Direct disruption of adipocytic identity regulators is insufficient to drive dedifferentiation

Because p300 functions as a central transcriptional co−activator and chromatin regulator, its loss is expected to destabilize multiple gene programs rather than act as a single lineage−specific driver of dedifferentiation. Nevertheless, the emergence of dedifferentiated regions upon *ep300* disruption therefore raised the question of whether dedifferentiation is mediated by loss of a specific downstream effector that directly enforces adipocytic identity. To address this, we focused on genes recurrently altered in dedifferentiated liposarcoma^3,8,9,16^ and with established roles in adipocyte differentiation, lipid homeostasis, or lineage stabilization^33–35^.

First, we investigated whether the disruption of an adipocyte maturation switch contributed to dedifferentiation in this model. To this end, we targeted *cebpa*, a key downstream effector of p300 during adipocyte maturation, where p300−dependent co−activation of C/EBPα is required to drive terminal adipocyte differentiation. Across liposarcoma subtypes, lower CEBPA expression resulting from DDLPS−specific 19q13 loss has been documented to be associated with poorer clinical outcome, linking impaired adipocytic differentiation programs to disease severity^8^. Targeting the master adipocytic differentiation regulator *cebpa* in our model in the same genetic background (i.e. **Δp53/Rb/Cebpa**) resulted in a limited incidence of low−grade well−differentiated liposarcoma (20%; n=8), while dedifferentiated lesions were not observed.

To test whether direct disruption of adipogenic identity contributed to liposarcoma progression, we replaced *ep300* targeting with loss of *g0s2* and *dgat2,* in the context of combined p53 and Rb pathway inactivation (**Δp53/Rb/adipo** cohort). In this setting, again no overt dedifferentiated liposarcomas were observed. Instead, approximately two−thirds (*i.e.* 69%, n = 16) of sampled adipose tissue exhibited cellular atypia, characterized by variability in cell size and shape without formation of bona fide lipoblasts or malignant architecture. These features are consistent with adipose dysplasia, resembling lipoma−like WDLPS, rather than high−grade (dedifferentiated) liposarcoma.

Finally, a **Δp53/Rb/Runx3** cohort was generated to test whether loss of the lineage−stabilizing factor Runx3, located on chromosome 1p, which is frequently deleted in DDLPS, is sufficient to promote liposarcoma dedifferentiation. Liposarcomas developed in only 13% of animals (n=31) and remained well−differentiated and did not display defined dedifferentiated regions.

Together, these experiments demonstrate that direct loss of adipogenic effectors or lineage−associated regulators induces adipocytic instability but is insufficient to drive malignant dedifferentiation. In contrast, dedifferentiation emerged only upon loss of upstream regulatory factors such as p300, indicating that disruption of higher−order transcriptional and chromatin−level control, rather than a combination of the above targeted genes, governs the dedifferentiated state in liposarcoma.

## Discussion

Retroperitoneal well-differentiated/dedifferentiated liposarcoma (WD/DDLPS) is characterized by extensive copy-number alterations, including recurrent gains and losses across the genome and, most prominently, amplification of the 12q13–15 region^17^. Within this region, co-amplification of MDM2 and CDK4 is routinely used for diagnosis and classification. In contrast, the broader amplicon spans many additional genes, and such large-scale alterations complicate efforts to resolve the specific genetic and regulatory events that drive tumor initiation, progression, and phenotypic heterogeneity. These alterations are widely used for diagnosis and classification, yet they typically encompass large genomic regions containing many genes, complicating efforts to resolve the specific genetic and regulatory events that drive tumor initiation, progression, and phenotypic heterogeneity. This challenge is particularly relevant for DDLPS, which exhibits increased cellularity, loss of adipocytic differentiation, and markedly worse clinical outcomes compared to WDLPS. Despite this biological and prognostic divergence, both phenotypes are largely managed as a single disease entity, and therapeutic strategies remain focused on shared amplification events rather than mechanisms underlying lineage destabilization and dedifferentiation. Consequently, the development of robust *in vivo* models that allow functional dissection of these processes has remained challenging, limiting translation of genomic insights into therapeutic strategies.

As with other cancers, sarcoma cell lines and patient-derived xenograft models have been developed and are widely used to characterize these tumors and identify therapeutic targets^36^. However these systems carry inherent limitations, including clonal selection, microenvironmental distortion, and limited control over tumor initiation. Conditional mouse models based on combined *Tp53* and *Pten* inactivation have more recently demonstrated that adipocytic sarcomas can be generated with high penetrance in a mammalian setting, yielding either predominantly mixed well- and dedifferentiated liposarcoma or a broader spectrum of liposarcoma subtypes depending on the tissue/cell line targeting strategy used^18,37^. These studies underscore both the value of conditional genetics and the importance of effective cell-of-origin targeting, but they rely on tissue-restricted Cre drivers that predefine the lineage context of tumor initiation. In contrast, we present a *Xenopus tropicalis* model that is cell-lineage enriched rather than tissue-restricted: multiplex CRISPR disruption of p53 and Rb signaling pathways is introduced mosaically in blastomeres giving rise to the cell lineage of interest, and tumor development depends on *in vivo* positive selection of true driver mutations. Importantly, the targeted gene set directly mirrors the core clinical driver logic of WD/DDLPS by functionally inactivating the p53 and Rb pathways, which in patients occurs downstream of *MDM2* and *CDK4* amplification. This combination of pathway-level clinical relevance and lineage-enriched selection provides a complementary *in vivo* framework to study liposarcoma initiation. Notably, our data further support that disruption of the p53 and RB pathways alone is sufficient to initiate well-differentiated liposarcoma, consistent with these alterations representing minimal clinical driver insults. This implies that other genes in the WD/DDLPS defining 12q13-15 amplified region, or additional recurrent amplifications frequently observed in patient tumors, are not essential for tumor initiation in this context. In contrast to existing mouse and zebrafish models, which often rely on genetic pathway perturbations not directly reflective of human WDLPS/DDLPS, our approach enables direct inference on clinically relevant minimal driver events.

A defining feature of human WD/DDLPS is the low burden of direct inactivating mutations in canonical cell-cycle tumor suppressors. *TP53* mutations occur in only a small fraction of cases (approximately 4% of DDLPS in TCGA), while *RB1* mutations are rare, and *RBL1* and *RBL2* are typically not mutated^4^. Instead, loss of p53–Rb pathway control is predominantly achieved through dosage-based mechanisms, most notably amplification of *MDM2* (≈97% of cases) and *CDK4* (up to ≈92% of cases). In contrast, these amplifications are far less frequent in non-WD/DDLPS sarcomas (≈19% and 4%, respectively)^38^. Together, these observations indicate that pathway disruption in liposarcoma is largely structural rather than mutational in nature, and that amplification of *MDM2* and *CDK4* may be sufficient to drive well-differentiated disease.

Interestingly, additional CRISPR-targeting of *ep300* destabilized adipocytic differentiation programs in our *Xenopus* model, resulting in increased phenotypic heterogeneity, with individual tumors containing spatially distinct regions that span a continuum from well-differentiated to non-lipogenic highly cellular dedifferentiated states. This intratumoral heterogeneity closely mirrors the clinical presentation of dedifferentiated liposarcoma, which is typically diagnosed as a mixed lesion containing both well-differentiated and dedifferentiated components^39,40^.

While several factors implicated in adipocytic programs, such as *CEBPA*, *G0S2*, *DGAT2* and *RUNX3* have been found to be lost in DDLPS^3,8,9,16^, targeted disruption of these individual components in the context of p53 and Rb deficiency did not phenocopy dedifferentiation in our model. Replacement of *ep300* targeting by *cebpa* or *runx3* loss resulted in slightly more malignant lesions compared to combined *g0s2*/*dgat2* targeting, yet neither condition gave rise to dedifferentiated tumors, in contrast to the robust DDLPS-like phenotype observed upon *ep300* loss. Notably, reduced CEBPA activity in DDLPS is frequently driven by epigenetic silencing rather than focal gene loss^41^, pointing to deregulation at the chromatin and transcriptional control level rather than isolated downstream factor deficiency. Together, these findings indicate that dedifferentiation is unlikely to result from loss of individual adipocytic regulators alone, but instead arises from disruption of higher-order regulatory mechanisms that maintain differentiation stability. In this context, *ep300* loss may act as a permissive event that destabilizes adipocytic identity at a regulatory level, not readily recapitulated by targeting downstream differentiation-associated genes. Together with the observed intralesional heterogeneity, this argues against a strictly linear progression from well-differentiated to dedifferentiated liposarcoma and supports a model of dynamically regulated adipocytic identity.

Together, these findings establish *Xenopus tropicalis* as an *in vivo* platform that recapitulates key molecular pathway disruptions and the defining histopathological heterogeneity of human retroperitoneal liposarcoma. Early embryonic mosaic targeting enables tumors to arise within permissive developmental contexts without imposing tissue-restricted assumptions, allowing direct *in vivo* interrogation of how genetic and regulatory perturbations shape tumor identity. This framework provides a foundation for resolving differentiation-state–specific dependencies in WD/DDLPS and may offer broader insights into plasticity-driven progression across other sarcoma subtypes.

## Material and Methods

### Generation of mosaic mutant animals and genotyping

Single guide RNAs (sgRNAs) were designed using CRISPRScan^42^ prioritizing target sites with high predicted on-target efficiency and minimal off-target risk. Candidate guides were further ranked based on predicted DNA repair outcomes following Cas9-induced double-strand breaks using the inDelphi algorithm, favoring sites with a high likelihood of frameshift-inducing indels, as previously validated in *Xenopus tropicalis* embryos^24^. Genome editing was performed by microinjection of sgRNA/Cas9 ribonucleoprotein complexes using recombinant NLS-Cas9-NLS protein^21^. All the CRISPR/Cas9 combinations described in the study were injected in one dorsal blastomere of four-cell stage *X. tropicalis* embryos and can be found in Supplementary Table 1, Sheet C. Injection mixtures were made by pooling sgRNA/Cas9 premixes that have been pre-incubated at 37°C for 30 sec. sgRNA dilutions for all the injection setups can be found in Supplementary Table 1, Sheet C. DNA from embryos (3dpf), tissues and tumors was extracted by means of an overnight 55°C incubation in lysis buffer (50 mM Tris pH 8.8, 1 mM EDTA, 0.5% Tween-20, and 200 μg/ml proteinase K) followed by heat inactivation. Amplicon deep sequencing and subsequent BATCH-GE analysis were performed as previously described^43^. All sgRNAs and primers used in this study are listed in Supplementary Table 1, Sheet C. All embryo, tissue and tumor sequencing data can be found in Supplementary Table 1, Sheet A-B.

### Phenotyping of cancer models

Microdissection images were taken using a Carl Zeiss StereoLUMAR.V12 stereomicroscope. Histological analysis was performed by means of classical H&E stains to get insights on tissue/tumor architecture and morphology. Tumor proliferation was assessed by immunohistochemistry using phospho-histone H3 (PHH3) and proliferating cell nuclear antigen (PCNA) as proliferation markers. Detailed information on primary and secondary antibodies and their respective concentrations is provided in Supplementary Table 1 (Sheet C). Chromogenic signal development was performed using the VECTASTAIN Elite ABC HRP Kit (PK-6100, Vector Laboratories) followed by ImmPACT DAB Peroxidase substrate (SK-4105, Vector Laboratories) according to the manufacturer’s instructions. For quantifications of cell numbers and tumor size areas the quPath quantification tool was used^44^. Histological images were acquired either via an Olympus BX51 Discussion Microscope or via a ZEISS Axioscan 7 machine at 20x magnification with a resolution of 0.22 μm/pixel. The pathological interpretations were made in compliance with a clinical pathologist (*i.e.* David Creytens, M.D.) at the University hospital in Ghent.

### Transplantation experiments

The generation of the immunocompromised *rag2^−/−^* line lacking the mature T and B cell pool was described before^26^. Tumor cell suspensions were generated by crushing tumor pieces with a pestle in 0.66x PBS (= amphibian PBS) and passing them through a 40 µm cell strainer (Falcon^TM^). Tumor cell suspensions were injected intraperitoneally or subcutaneously at the back of the animals using BD Micro-Fine Demi 0.3 mL 0.3 mm (30G) x 8 mm syringes. Transplanted animals were closely monitored and euthanized when showing early symptoms of discomfort. All animal experiments were conducted in accordance with local legal and institutional guidelines and approved by the ethical committee of the Center for Inflammation Research and Ghent University, Faculty of Sciences.

### Statistical analyses

All statistical analyses were performed by means of two-sided student’s t-tests, with Welch-correction (when homoscedasticity criterium was violated) and can be found in Sup. Table 2. Significance intervals are shown in the figures and are as followed: non-significant *p* ≥ 0.05, \**p* < 0.05, \*\**p* < 0.01. Graphs are represented as population means with error bars illustrating S.E.M. values.

## Acknowledgements

Research in the authors’ laboratory is supported by the Research Foundation—Flanders (FWO-Vlaanderen) (3G0A6922), by the Foundation against Cancer (365L07823), and by the Concerted Research Actions from Ghent University. David Creytens was financially supported by a senior clinical research fellowship from the Research Foundation Flanders (FWO-Vlaanderen) (1800725N). We would like to acknowledge Benjamin Pavie (VIB Bioimaging Core) for assisting in the QuPath quantifications. Furthermore, we are thankful to Tim Deceuninck for the good animal care.

## Contributions

M.B., D.T. and K.V. designed the study. M.B. and D.T. performed all CRISPR–Cas9 injections. M.B., D.T., A.C., W.H. and S.D. conducted genotyping and phenotyping experiments. D.T. performed the transplantation experiments. D.C. carried out the pathological assessment. M.B. and D.T. analyzed the data and generated the figures. M.B. and D.T. wrote the manuscript. K.V. and D.C. revised the manuscript.

## Supplementary Figures

**Supplementary Figure 1.**
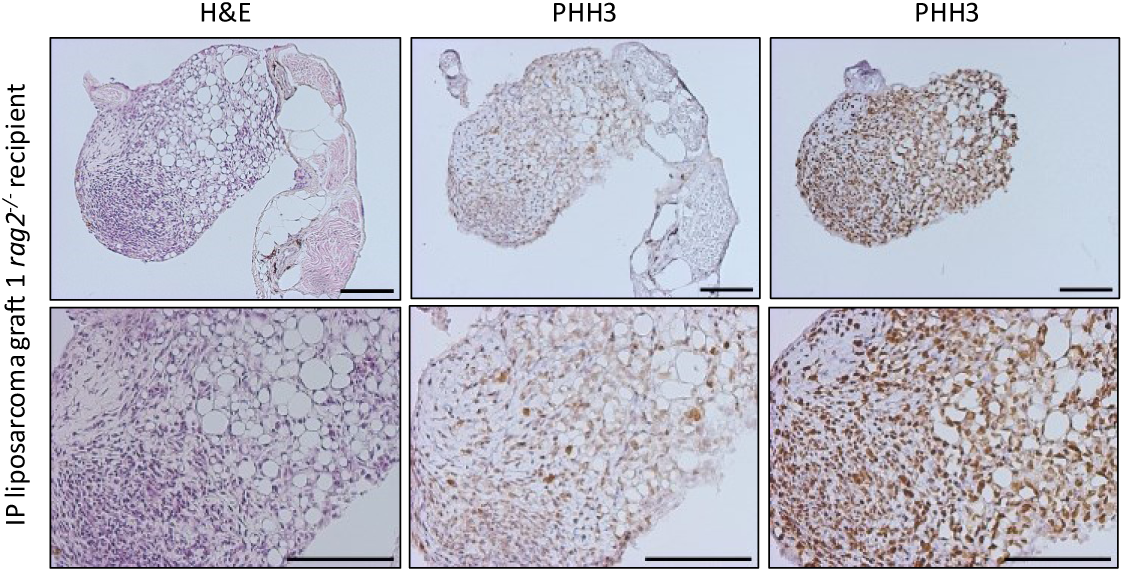
Proliferation analysis of Δp53/Rb/p300-derived liposarcoma graft in immunodeficient rag2^−/−^ Xenopus tropicalis. Intraperitoneally (IP) transplanted donor-derived liposarcoma grafts in rag2⁻/⁻ recipients display high proliferative activity, as evidenced by positive staining for the proliferation markers PHH3 and PCNA. Scale bars: 100 µm

**Supplementary Figure 2.**
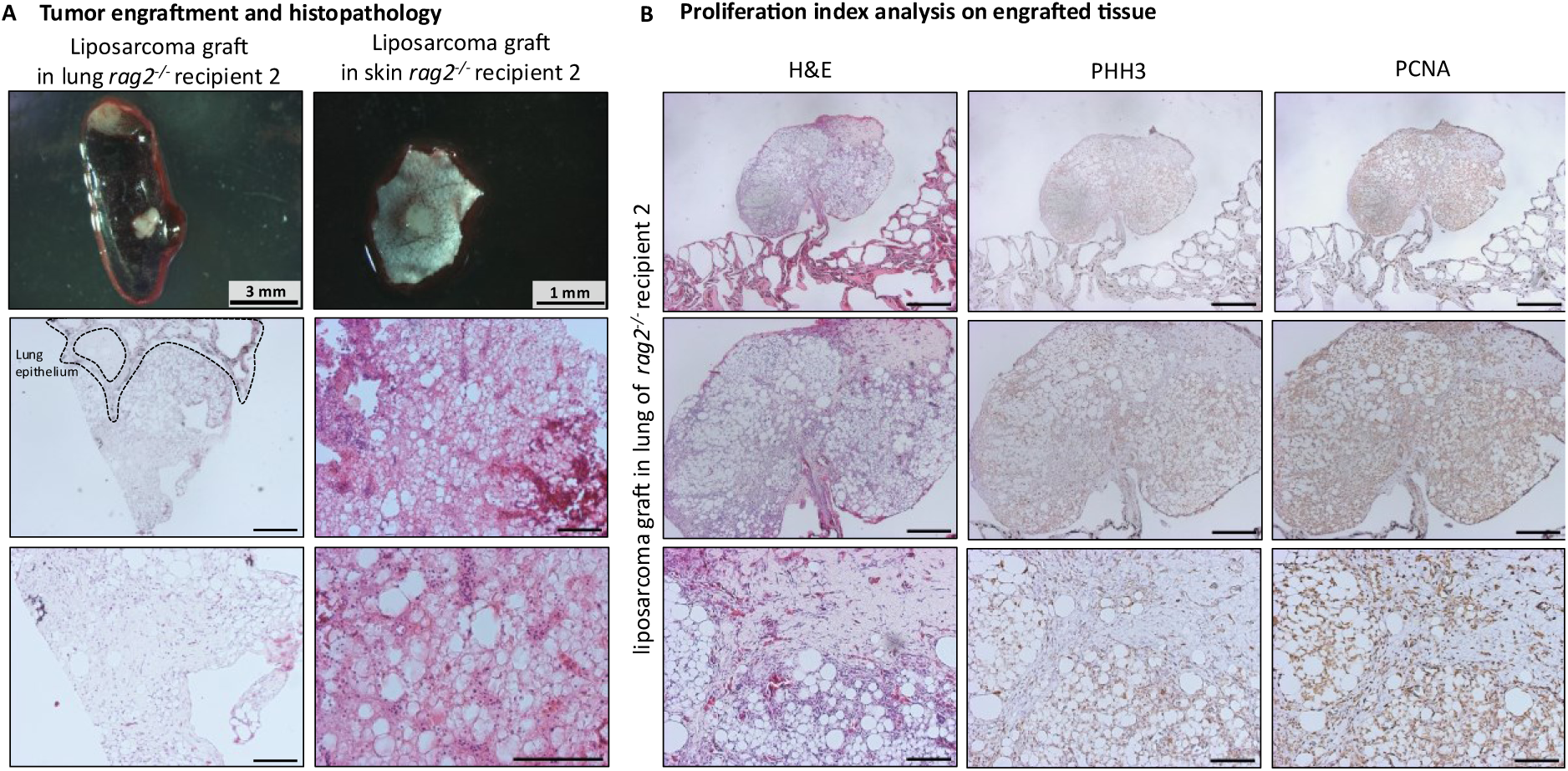
Parallel allotransplantation of Δp53/Rb/p300 liposarcomas in immunodeficient rag2^−/−^ Xenopus tropicalis. **(A)** Independent confirmation of tumor engraftment is shown in a second rag2^⁻/⁻^ recipient, with donor-derived liposarcoma cells detected in lung tissue and at the intraperitoneal (IP) injection site by histopathology. **(B)** The grafts also display high proliferative activity, as evidenced by positive staining for the proliferation markers PHH3 and PCNA. Scale bars histology: 500 µm and 100 µm (middle row), 100 µm (bottom row). Scale bars IHC: 500 µm (top row), 250 µm (middle row), 100 µm (bottom row).

**Supplementary Figure 3.**
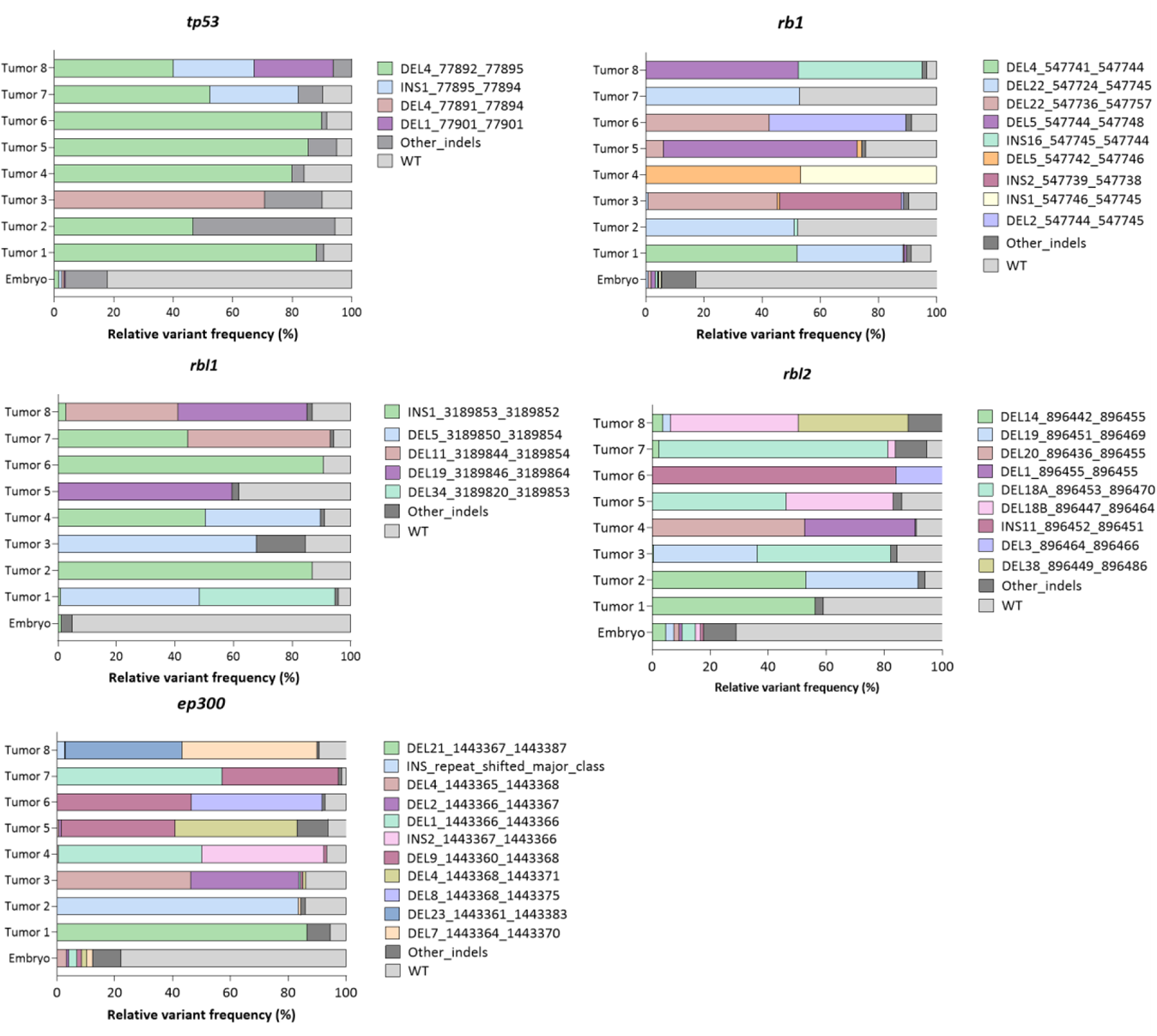
Selection of dominant CRISPR-induced indel variants in liposarcoma tumors. Stacked bar plots showing the relative frequencies of indel variants for five targeted genes (tp53, rb1, rbl1, rbl2, ep300) in whole-embryo samples and eight tumors (tumor 1 – tumor 8). Each bar represents one sample; colors denote specific variants, with remaining low-frequency mutations grouped as Other indels and wild-type (WT) reads shown. Whole-embryo samples display diverse, low-frequency editing, whereas tumors are dominated by one or a few variants, indicating clonal selection.

**Supplementary Figure 4.**
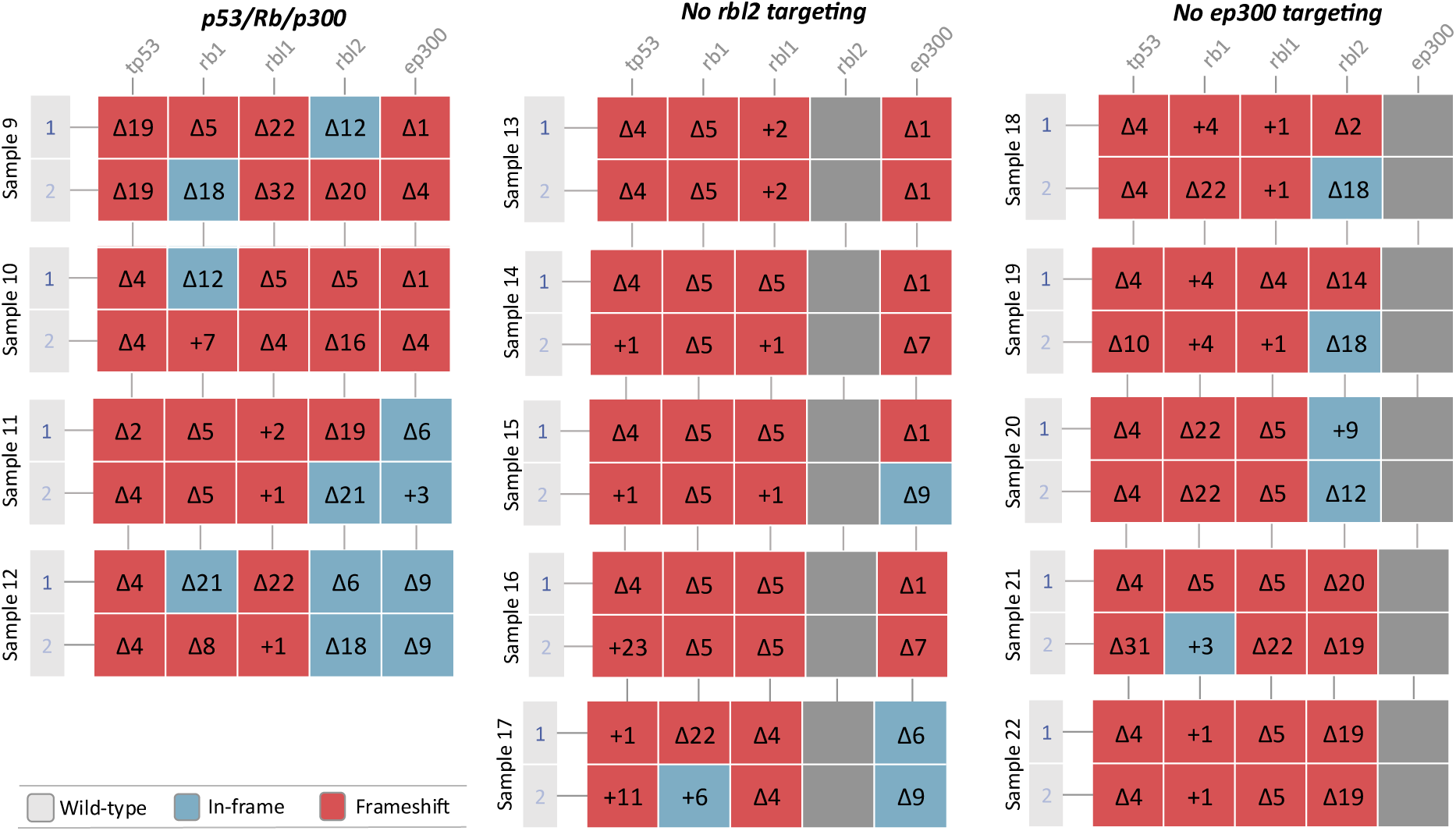
Additional deep amplicon sequencing confirms positive selection on biallelic mutated gene status in Δp53/Rb/p300 liposarcomas. Allele-specific mutation status across fourteen additional liposarcoma samples of Δp53/Rb/p300, Δp53/Rb(-Rbl2), Δp53/Rb cohorts. Two rows per sample represent inferred alleles. Colors indicate wild-type (grey), in-frame (blue) and frameshift (red) indels.

**Supplementary Figure 5.**
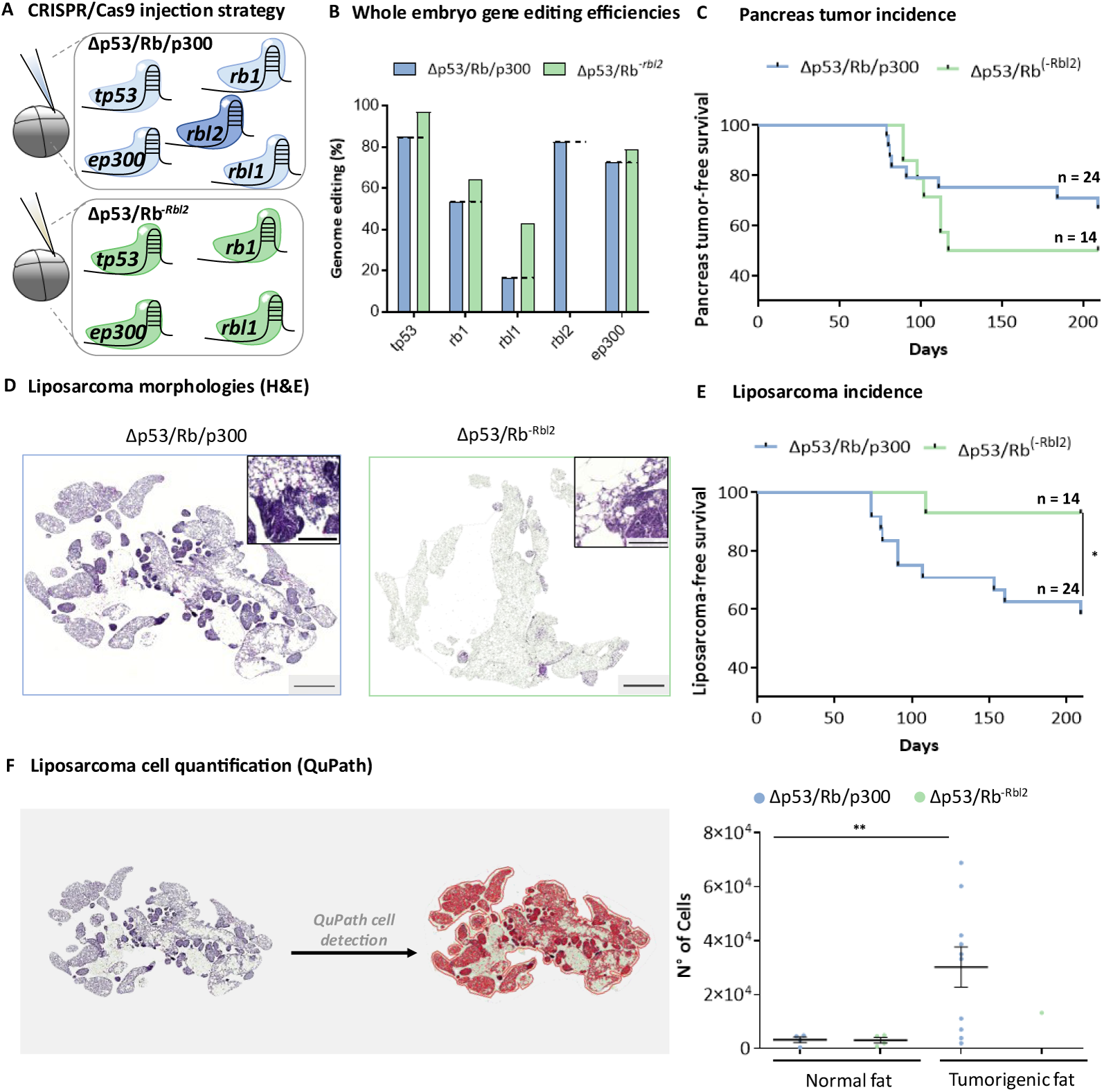
Loss of rbl2 drives liposarcoma dedifferentiation. **(A)** Experimental design comparing multiplex CRISPR targeting of Δp53/Rb/p300 with or without ep300 disruption. **(B)** Estimated whole-embryo genome editing efficiencies for each targeted gene across injection cohorts. Total efficiencies were extrapolated by multiplying measured efficiencies by four, as embryos were injected in one of four blastomeres. Values exceeding 100%—likely due to incomplete cytokinesis allowing reagent diffusion to the sister blastomere—were normalized such that the most efficient sgRNA was set to 100%. **(C)** Pancreas tumor-free survival compared between Δp53/Rb/p300 and Δp53/Rb^−Rbl2^ cohorts. **(D)** Representative H&E-stained sections illustrating liposarcoma morphology across Δp53/Rb/p300 and Δp53/Rb^−Rbl2^ conditions. **(E)** Liposarcoma-free survival analysis comparing Δp53/Rb/p300 and Δp53/Rb^−Rbl2^ cohorts. Scale bars histology: 500 μm and 100 μm (inset). **(F)** Quantification of liposarcoma cell numbers using QuPath. For each animal, the largest tumor section was analyzed (representative image shown on the left, all are listed in Supplementary Fig. 5). Four representative healthy littermate adipose tissue samples were included as controls to establish baseline adipocyte cellularity.

**Supplementary Figure 6.**
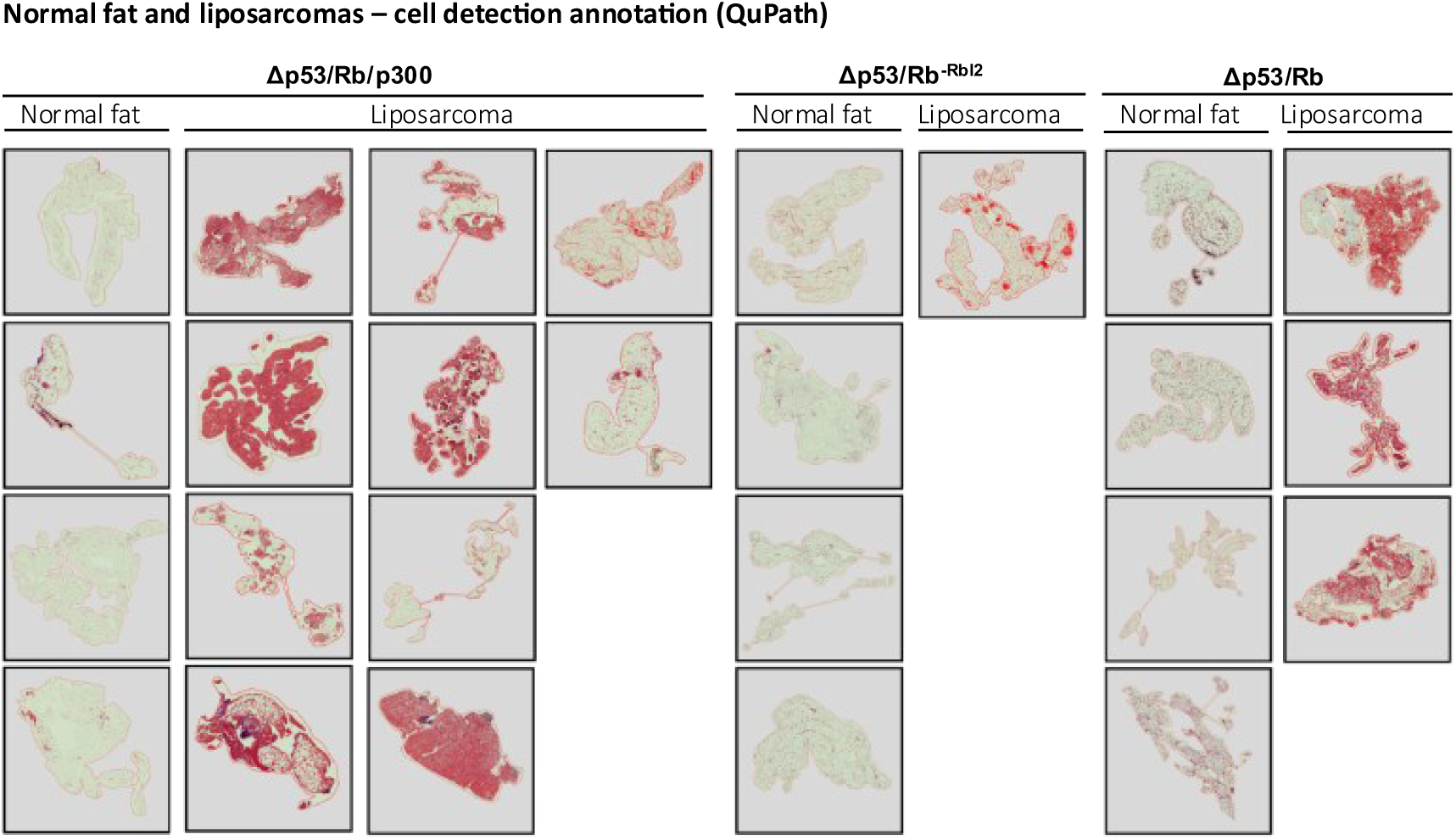
QuPath quantification algorithm reveals differences in tumor characteristics in Δp53/Rb/p300, Δp53/Rb^−Rbl2^ and Δp53/Rb liposarcomas. Overview of all liposarcoma versus normal fat sections (n = 4) for all liposarcomas sampled in the Δp53/Rb/p300 (n = 10), Δp53/Rb^−Rbl2^ (n = 1) and Δp53/Rb (n = 3) cohorts (sections with largest respective tissues used as input), quantified with the QuPath software and calculated as cell numbers. Each hematoxylin cell is detected, pseudo-stained in red and counted with the software.

## Supplementary Tables

Supplementary Tables are provided as separate Excel files. Supplementary Table 1 (Sheets A–C) includes PCR-based genotyping, amplicon deep sequencing (MiSeq), BATCH-GE analysis, and DNA oligo and antibody information. Supplementary Table 2 (Sheets A–F) contains all statistical analyses.

